# Integrative Molecular Dynamics Simulations Untangle Cross-Linking Data to Unveil Mitochondrial Protein Distributions

**DOI:** 10.1101/2024.09.11.612425

**Authors:** Fabian Schuhmann, Kerem Can Akkaya, Dmytro Puchkov, Martin Lehmann, Fan Liu, Weria Pezeshkian

**Author notes:** These authors contributed equally (Fabian Schuhmann, Kerem Can Akkaya).

## Abstract

Cross-linking mass spectrometry (XL-MS) enables the mapping of protein-protein interactions on the cellular level. When applied to all compartments of mitochondria, the sheer number of cross-links and connections can be overwhelming, rendering simple cluster analyses convoluted and uninformative. To address this limitation, we integrate the XL-MS data, 3D electron microscopy data, and localization annotations with a supra coarse-grained molecular dynamics simulation to sort all data, making clusters more accessible and interpretable. In the context of mitochondria, this method, through a total of 6.9 milliseconds of simulations, successfully identifies known, suggests unknown protein clusters, and reveals the distribution of inner mitochondrial membrane proteins allowing a more precise localization within compartments. Our integrative approach suggests, that two so-far ambigiously placed proteins FAM162A and TMEM126A are localized in the cristae, which is validated through super resolution microscopy. Together, this demonstrates the strong potential of the presented approach.

**Entry for the Table of Contents.**
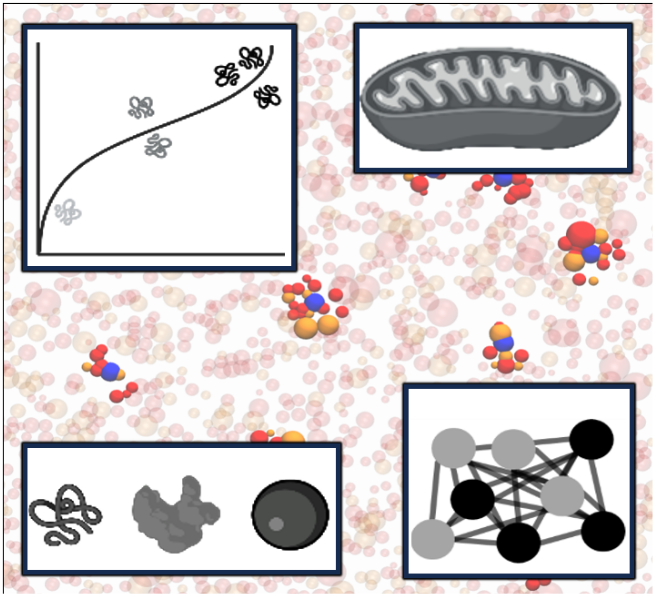
The approach combines cross-linking mass spectrometry, electron microscopy, and coarse-grained molecular dynamics simulations to map protein-protein interactions in mitochondria. This integration makes complex data clusters accessible, revealing protein distributions and localizations. Notably, it allows the localization of FAM162A and TMEM126A through super-resolution microscopy, showcasing the synergy of experimental and computational methods.

## Introduction

Protein-protein interactions and the resulting formation of molecular complexes and protein clusters are fundamental to the functional integrity and regulation of cellular and organellar processes. ^[1]^ Such interactions are abundant, subtle and often poorly or not at all understood. Since characterizing protein-protein interaction networks is essential for understanding the functional mechanisms of cells and organelles; however, this task remains a significant challenge.

Numerous efforts and approaches are undertaken throughout the scientific community to unravel interaction patterns, which led to the discovery of hundreds of thousands of human protein interactions. ^[2–4]^ Some of these techniques include proximity biotinylation, affinity purification mass spectrometry, and native mass spectrometry, each contributing uniquely to mapping the cellular interactome. ^[5–9]^ Another one of these experimental tools to identify the protein interactions is cross-linking mass spectrometry (XL-MS). ^[10]^ XL-MS utilizes chemical cross linkers (a linker with two functional groups and an adjustable spacer) which selectively bind to specific amino acids, linking two proteins together.^[11]^ As such, cross-linking can only occur, if the proteins were close enough to each other. ^[12]^ Therefore, XL-MS enables a detailed characterization of the protein-protein interactions within a predefined radius, correlating to the closeness of the proteins in the sample and their likelihood of having formed protein complexes.^[13–17]^

However, employing XL-MS experiments to quantify interactions in larger systems, i.e. cells and organelles, results in convoluted and inconclusive networks of the protein interactions.^[18–20]^ Despite the development of proteome-wide XL-MS methods, which have demonstrated the ability to capture extensive portions of the proteome in intact cells and organelles, they have been mostly utilized for analyzing protein structures and interactions. ^[21–23]^ However, recently, Zhu *et al*. ^[17]^ characterized the interaction network in mitochondria to extract protein localization and extend the mitochondrial proteins’ annotation. The resulting interaction network shows very large clusters (on the magnitude of hundreds of proteins). However, the known protein complex sizes in mitochondria contain just up to 50 individual sub-units which are lost in the network. ^[24,25]^ A visualization of the clustering is shown in Fig. 1.

**Figure 1.**
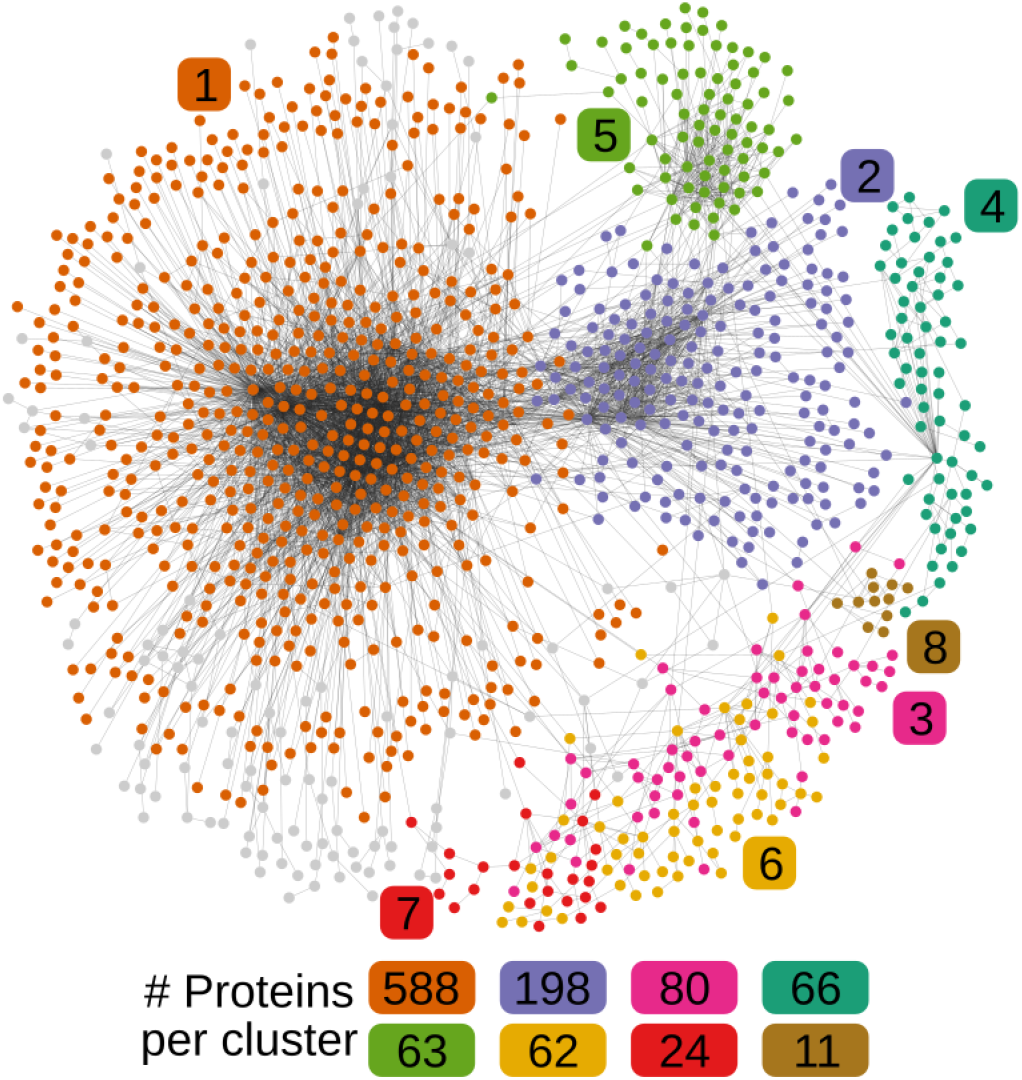
Visualization of edge betweenness cluster analysis on XL-MS data obtained from Zhu *et al*. ^[17]^. Differently colored nodes show their major cluster assignment (>10 members), while grey nodes remain unassigned. The colored numbers indicate the number of proteins in the corresponding clusters. An edge-weighted spring embedded layout was applied based on cluster assignment. Visualization performed using Cytoscape.^[26]^

As the clusters do not paint a clear picture, further distorted by varying protein abundance levels, our understanding of the spatial arrangements, the interactome, the assembly of macromolecular complexes, and the localization of individual protein types is still vague at best.

To remedy this problem, endevours to interface experiments with *in silico* studies are undertaken. ^[27]^ We employ an supra coarse-grained integrative molecular dynamics model (MD), calibrated using crosslinking mass spectrometry (XL-MS) data, quantitative mass spectrometry, and 3D volumetric imaging techniques. Through the here presented model, we obtain insight into protein distribution and cluster formation in a experimentally backed-up interaction strength through reasonable cluster sizes. The MD simulations recovers known clusters from the data, which validates the method. At the same time, previously unknown clusters are highlighted which inspires optimism in yet-to-be discovered protein clusters. Additionally, we provide an overview of the distributions of a majority of mitochondrial proteins, which can be used to infer the localization of certain proteins in the different mitochondrial compartments of the, e.g. inner mitochondiral membrane. The localization is then further verified through stimulated emission depletion (STED) microscopy.

## Results and Discussion

### Cluster analysis of XL-MS data is inconclusive

For this study, only mitochondrial proteins were investigated to distinguish the functional protein clusters and distributions within the organelle. With a total of 1326 proteins found in the previously published XL-MS data ^[17]^, the data allows the generation of a matrix counting the number of cross-links between the protein types, pairwise. The resulting matrix is a symmetric matrix with dimensions 1326 *×* 1326 with 20652 entries not being zero. The XL-MS experiment thus finds 10326 pairs of different proteins, which share at least one cross-link (see Fig. 1). Such a matrix can be drawn as a graph network and clustering analysis can be performed. Figure 1 shows such a network with clusters that contain more than ten proteins colored. Gray proteins are unclustered. Further details on the utilized experimental data can be found in the supplementary information S1.

The dataset contains 1326 proteins from purified mitochondria. In the purification process, associated membranes are also obtained, which adds cytosolic proteins to the total number, which are excluded from the present study to focus on mitochondrial proteins. A total of 713 different proteins belong to the different mitochondrial compartments, of which 608 have a measured protein abundance.^[28,29]^

The clustering isolates eight clusters with sizes ranging from 11 proteins in the smallest cluster to 588 proteins in the largest cluster. As the protein cluster sizes, for, e.g., the respiratory complexes ranges from 4 to 45 protein units, these clusters are lost in the analysis. Hence, it was only analyzed whether a protein is mitochondrial or not, and if so, to which compartment the protein might belong.

### An integrative supra coarse-grained model is built

We propose a stochastic dynamics simulation, which manifests as a variation to a classical mechanics molecular dynamics simulation. The simulation is conducted in a supra coarse-grained method, in which each protein is modeled by a single particle and the following assumptions are made.

- The XL-MS data accounts for all the interactions between the different protein types.
- Each protein’s size can be approximated through its radius of gyration.
- Proteins will not form covalently-bonded multimers.
- Protein abundancy is normalized to the simulated volume for each compartment.

The technical aspects of the simulation are described in the supplementary information S2.

The interactions between two proteins *A* and *B* which have a distance of *rAB* is governed by a Lennard-Jones (LJ) type potential. The LJ potential is typically used to describe the van der Waals interaction in an atomistic simulation ^[30,31]^, which simply includes attractive and repulsive forces and can easily be modulated by the interactions obtained from the experimental data. The modulation is done proportional to the number of cross links between the different protein types. Hence, the interaction is given by

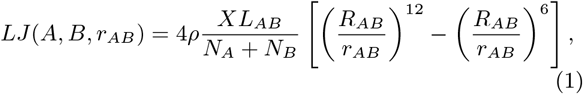

with *XL*_*AB*_ describing the number of cross-links between protein *A* and *B, NA* and *NB* are the abundances of the proteins as obtained by Morgenstern *et al*. ^[29]^. *R*_*AB*_ is the arithmetic mean of the radius of gyration of proteins *A* and *B*. The scaling factor *ρ* scales the value to reasonable magnitudes (*ρ* = 60), which was obtained through a calibration process only considering the OMM compartment, which is described in detail in the supplementary information S3. The third assumption guarantees, that no bonded interactions have to be considered in the simulation. The first assumption thus reduces all interaction terms to the Lennard-Jones potential. Equation (1) therefore describes an effective potential including all interactions between two proteins, e.g, Van der Waals, electrostatic and even hydrophobic driven forces. Additionally, the intrinsic curvature of the membranes and their influence on protein dynamics is not explicitly captured in the MD simulation, the curvature influenced interactions are in fact implicitly in the XL-MS data.^[32]^ The radius of gyration was approximated by the radius obtained from AlphaFold2 ^[33]^ structures based on the sequence of all the encountered proteins.

The fourth assumption requires the determination of individual protein abudances scaled to the volume of the simulation. These abundances are estimated through 3D volumetric electron microscopy (EM), which is described in detail below. In total, some proteins might be too rare to have a single copy in the simulation box, thus reducing the number of simulated proteins to 491.

Figure 2A-D pictographically visualize the different data types and experiments which inform the MD simulation, while Fig. 2E shows a cross section of mitochondria that serves as a model for the compartments in the simulation, chosen for including all mitochondrial compartments. Figure 2F visualizes the intial situation of the MD simulation from which the interactions are computed. Proteins are sorted into their respective compartments based on the data obtained from MitoCarta 3.0. ^[28]^ The compartments are denoted as follows:

- OMM: Outer Mitochondrial Membrane
- IMS: Intermembrane Space
- IMM: Inner Mitochondrial Membrane
  - Cristae: Cristae
  - IBM: Inner Boundary Membrane
- Matrix: Matrix

**Figure 2.**
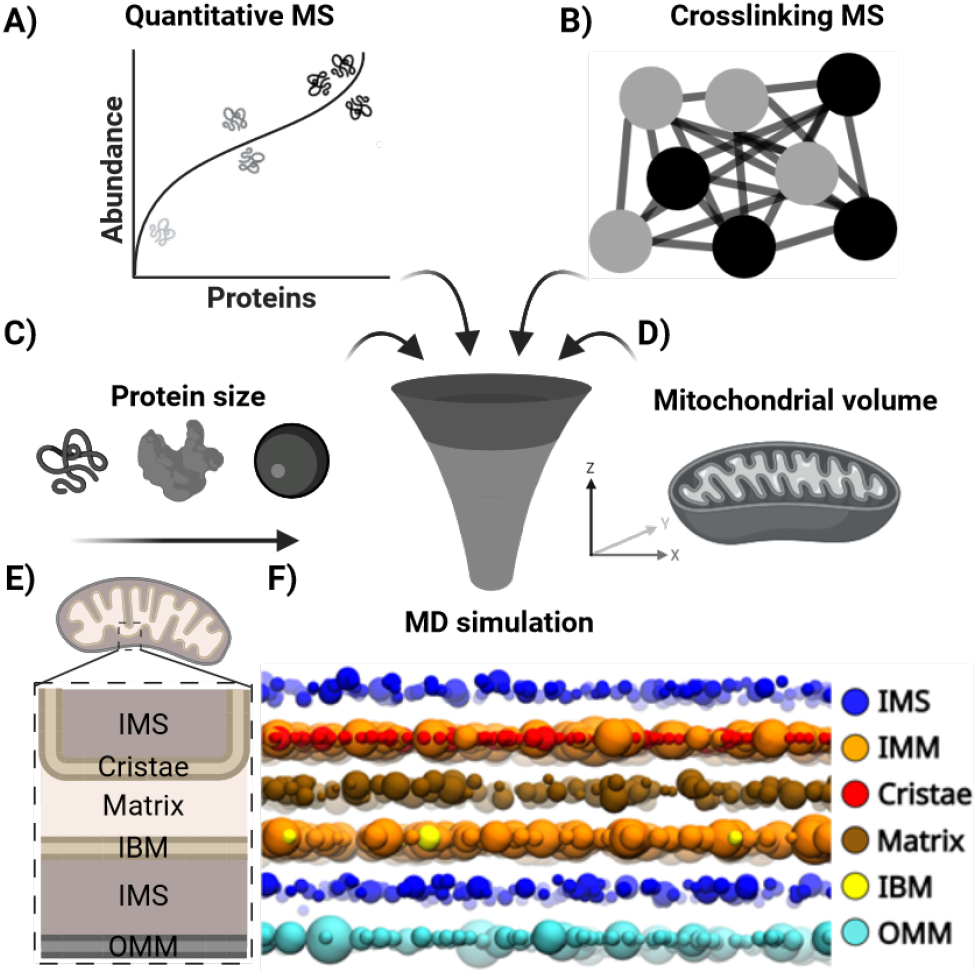
A) Quantitative MS data regarding mitochondrial protein copy numbers per cell were obtained from Morgenstern *et al*. ^[29]^. B) Cross-linking MS data on mitochondria were obtained from Zhu *et al*. ^[17]^. The DSBSO dataset was utilized for analysis. The mass spectrometry raw data has been obtained from the PRIDE repository with the dataset identifier PXD046382. C) The proteins are represented as globular balls, based on the radius of gyration as obtained from the predicted AF2 structures. D) Mitochondrial volumetric data was acquired by combining light microscopy and 3D electron microscopy. Experimental details can be found in the supplementary information S1. E) Schematic model of the section of the mitochondrion that inspired the MD simulation setup. The section was chosen because it is a generic section that includes all compartments. F) Initial protein setup for the MD simulation. Proteins are colored according to their assigned compartment. Visualization is rendered with VMD. ^[38]^ (Panels A-E were created with BioRender.com.)

Proteins located in the cristae or the IBM are also part of the IMM, but their localization along the membrane to the different regions is known. ^[34,35]^ The other proteins in the IMM cannot be directly attributed to a certain area and are thus placed randomly in both membranes. ^[36,37]^ In the simulation, membrane proteins are constrained to only move in a plane that symbolizes a flat membrane, while the soluble proteins (IMS, Matrix) are free to move between membranes. They cannot cross membranes, however.

### Volumetric data is needed to complete the simulation setup

We used a microscopy-based approach to determine the MD simulation space and normalize protein abundances to determine the absolute and relative total mitochondrial volume relative to the total cell volume. This approach, visualized in Fig. 3, combines confocal microscopy of living cells and electron microscopy of fixed cells as orthogonal controls. Under standard cultivation conditions, the cellular volume of HEK293T was measured as roughly 5, 400 *µm*^3^. The total volume of all mitochondria was measured at an average of 543 *µm*^3^, representing roughly 10% of the total cellular volume (Fig 3A, D). As our model employs protein abundances for the whole cell and, hence, all the cell’s mitochondria, these abundances need to be scaled for a single mitochondrion and, respectively, for the volumina of different compartments.

**Figure 3.**
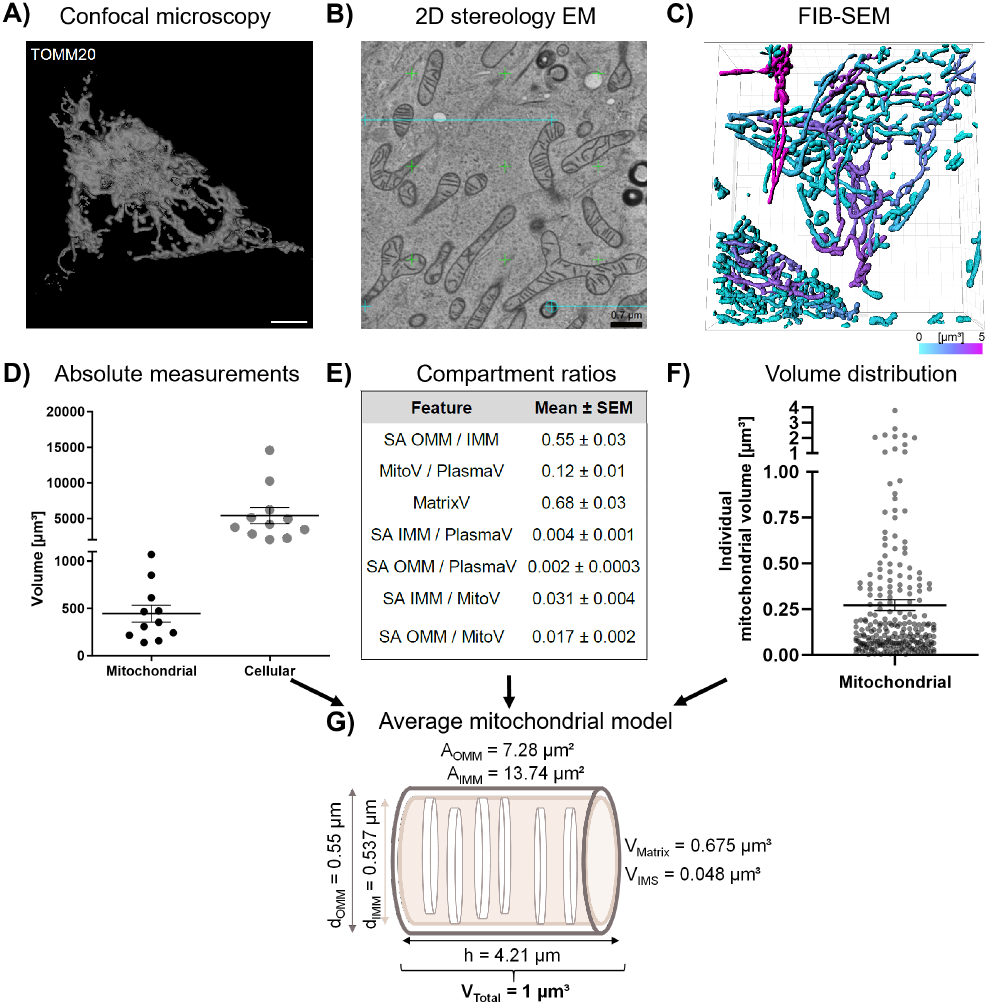
A) Maximum intensity-based 3D projection of acquired confocal z-stacks of single HEK293T cells stained for TOMM20, visualized in 3D Viewer. Manual selected thresholding was applied to match confocal image structures, resampling was set to 2. Scale bar: 10 *µm*. B) 2D stereology-based grid (green & cyan) on top of FIB-SEM sampled images. Scale bar: 0.7 *µm* C) 3D segmentation of mitochondria from FIB-SEM acquired z-stacks for analysis of individual mitochondrial volume. Mitochondria are divided into separate units of various volumes and colored according to their size on a gradient scale of 0 *µm*^3^ (cyan) up to 5 *µm*^3^ (magenta). Scale bar: 5 *µm*. D) Quantification of the mitochondrial and cellular volume based on absolute volume measurements of the 3D volumetric models using Imaris™. Single, separate cells were selected for quantification. *N* = 11 different cells. E) Relative ratio calculations from 2D stereology analysis for different features. The mean with the SEM is shown. MitoV = mitochondrial volume, PlasmaV = volume of the cytoplasm, MatrixV = matrix volume, SA = surface area. *N* = 22. F) Quantification of individual mitochondrial volumes, acquired from 3D segmentation from Panel C. Analysis is based on absolute volume measurements of the 3D volumetric models using Imaris. *N* = 231 measured mitochondria. The mean with the SEM is plotted. G) Abstraction of mitochondrial model as a cylinder with defined volume of 1 *µm*^3^ and diameter for extraction of defined surfaces and compartment sizes. Dark brown cylinder represents OMM, light brown cylinder represents IMM, with cross-sections representing cristae.

To obtain these nanoscale dimensions of surfaces and volumes of mitochondria and their subcompartments, stereology measurements were applied on 2D electron microscopy images of cross sections of HEK293T cell. The relation of the total volume fraction of mitochondria to its compartments was estimated (Fig. 3B, E), which included the volume spaces and their corresponding membrane compartments. Thus, matrix volume was determined to be around 68% of the total mitochondrial volume. At the same time, the IMM membrane surface was measured to be twice as big as the OMM. Consisting of the IBM and cristae area, this IMM area totaled to roughly 0.03 *µm*^2^, while the OMM surface area was measured to be around 0.016 *µm*^2^ per 1 *µm*^3^ of the mitochondrial volume.

The 3D FIB-SEM imaging and reconstruction showed that the volume of individual mitochondria spans from 0.27 *µm*^3^ to as high as 3.8 *µm*^3^ (Fig. 3C,F) It corresponds to live imaging data that show that mitochondria are highly dynamic organelles that can change their shape and undergo constant fission and fusion cycles depending on the different cellular states, either leaning to form interconnected, tubular networks or disintegrate into smaller individual mitochondrial objects. Since mitochondria show very diverse lengths and volumina, instead of trying to calculate an average individual mitochondria, a cylindrical mitochondrial model with a total mitochondrial volume of 1 *µm*^3^ was employed. The compartments were then built with the area and volumetric values obtained from the stereology approach to define absolute volumes for the MD simulation (Fig. 3G) Using the measurements and ratios from this cylindrical model, each membrane is represented in the MD simulation as a 2D 500 *×* 500 *nm*^2^ flat dimension. The soluble compartments are represented as 3D volumes with a defined height of 14*nm*. This leads to a simulation volume of 500 *×* 500 *×* 42*nm*^3^.

### Simulation results allow a refined cluster analysis

With this model, 30 simulations were run for 230 *µ*s each, starting from a random placement of all proteins in their respective compartments. The last simulation snapshot of each simulation was then taken to extract the neighbors of each protein type. Two proteins *A* and *B* were considered neighbors, if *r_AB_ <* 1.25 *· R_AB_*. The factor 1.25 is chosen to slightly extend the distance at which two proteins are considered close, because at *r_AB_* = *R_AB_*, the strong repulsion from the Lennard-Jones potential, would result in a negligible number of neighbors. The additional 25% results in an average neighbor radius of 0.79 *±* 0.17 *nm* (error standard deviation). Thermal fluctuations move attractive particles in our system beyond this range.

For each protein type, it was counted how many neighbors of another protein type were found and scaled by the abundance of the protein. This calculation yields a 491*×*491 matrix with entrys describing the relationship between the different protein types. The full matrix is available in the supplementary information S3. Then a cluster analysis was performed on the matrix to probe for known and unknown clusters alike.

The analysis show that the simulations predict some of the known cluster. For instance, in the clusters, the respiratory complexes I-V can be recovered. Also, the Pyruvate dehydrogenase (PDH) complex, parts of the translocase of the outer membrane (TOM) complex, and the detailed interaction landscape of molecular chaperones, i.e., with the designated gene names PHB1, PHB2, HSPA9, or TRIAP1. A detailed table of all measured neighborhoods from the MD simulation can be found in the supplementary information S4. Additionally, reduced matrices containing the protein neighborhoods from the MD simulation for the respiratory complexes can be found in S5. The standard deviation for the measured neighbors can be found in S6.

Expanding on the direct neighbor relationship between the protein types in the recovered interaction matrix, the simulation resolves the extensive interaction landscape within mitochondria (Fig. 4A) and a comparison to crosslinking derived data (Fig. 4B). Of the 491 proteins present, 229 are from the matrix, where they exhibit a majority of the interactions among other matrix proteins as well as with proteins of the IMM and the cristae. Only a few matrix to IBM interactions are observed. This can be attributed to the facts, that the abundance of IBM proteins is low and that not many proteins can be assigned to the IBM. The 42 OMM proteins mostly formed interactions within their compartment, but a few interactions could be observed to IMS proteins. The manually assigned 58 cristae proteins exhibited mainly interactions with IMM and matrix proteins, while the 15 IBM proteins showed a few separate interactions to all admissable compartments (IMM, matrix, IMS). The remaining unassigned 114 IMM proteins interacted mostly with cristae or matrix proteins and only with a few IBM interactions. MitoCarta 3.0 ^[28]^ assigns these proteins to the IMM and does not provide further information on their sub-compartmental distribution to the cristae or IBM. We investigated the distribution of IMM proteins by combining the MD simulation’s interaction data together with localization markers, similar to the cross-link derived CLASP approach employed by Zhu *et al*. ^[17]^; the CLASP approach employs interactions to known proteins and their compartmental localization to derive the localization of unknown proteins. Whereas CLASP is crosslinking based and restricted to the major compartments, we utilized the CLASP principle for elucidating the IMM protein distribution. Based on this, IMM proteins solely interacting with cristae proteins would be assigned to the CM. Out of the 114 IMM proteins present in the simulation, 77 proteins possess interactions to both cristae and IBM, with the rest only showing interactions to cristae. To further distinguish compartments, we introduced a ratio of the cristae and IBM interactions, visualized in Fig. 5A.

**Figure 4.**
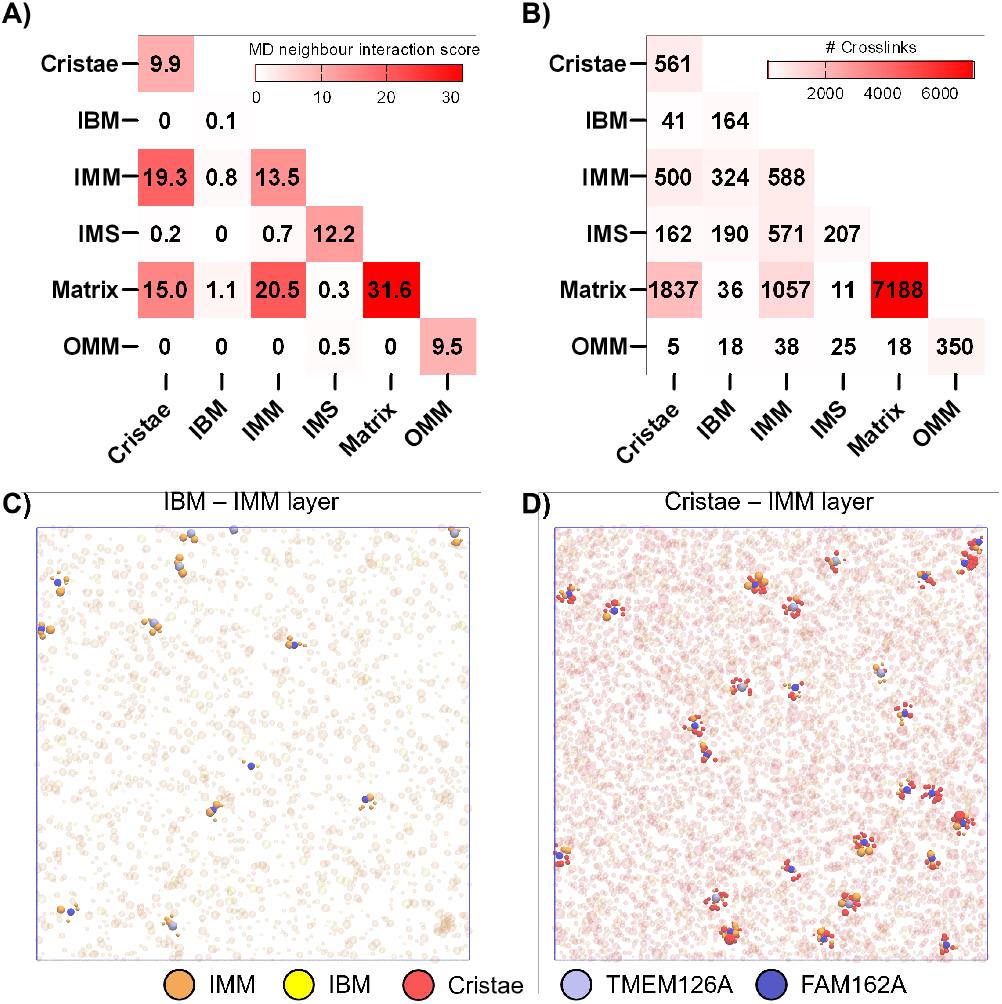
A-B) Distribution of observed MD (A) and XL-MS interactions (B) across the subcompartments and between different compartments. For the MD simulation, values are shown as the sum over all proteins in each compartment and all simulation replicas. For the XL-MS data, the number of unique crosslinks is shown. Sub-mitochondrial annotation is based on MitoCarta3.0 for the major compartments and based on literature evidence for selected IBM and cristae proteins as localization markers. C-D) The TMEM126A (light blue) and FAM162A (blue) proteins from the IMM compartment are shown in the IBM-IMM layer (C) and the Cristae-IMM layer (D). For visualization purposes the radius of TMEM126A and FAM162A has been doubled. Other proteins from the layer are high-lighted within 12.5 *nm* of the proteins. One notices that hardly any IBM proteins are located in the vicinity of the proteins, but cristae proteins can be found in abundance.

**Figure 5.**
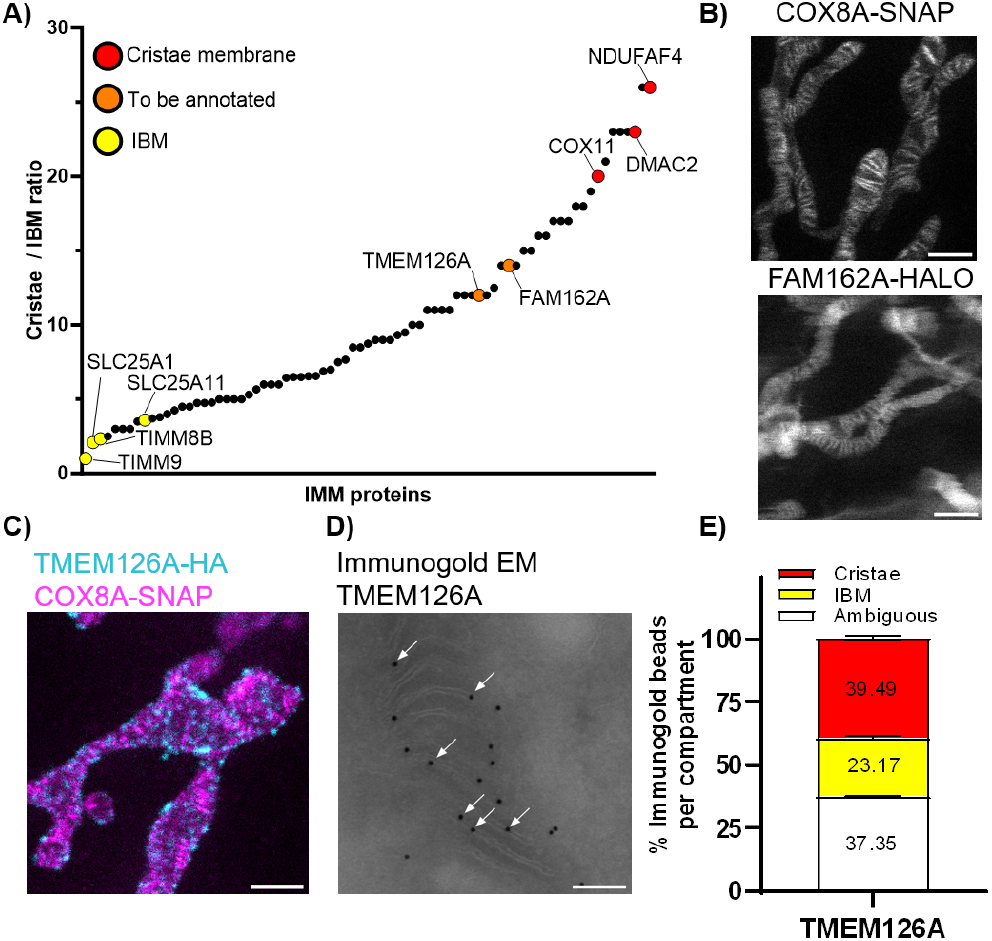
A) Distribution of Cristae / IBM interaction ratio for unassigned IMM proteins. Proteins are colored according to their suspected localization within the IMM: Cristae (red), IBM (yellow) and selected candidates (orange). B) Live STED imaging of COX8A and FAM162A. For live STED microscopy, COX8A-SNAP and FAM162A-Halo are stained using BG-SiR-d12 and CA-SiR-d12 in transfected HeLa-COX8A-SNAP cells. Scale bar: 1 *µm*.^[45]^ C) Fixed STED imaging of TMEM126A as a selected candidate. IMM(magenta) is stained in fixed HeLa-Cox8-SNAP cells using BG-SiR-d12 and HA-antibody against TMEM126A-HA (cyan). Scale bar: 1 *µm*. D) Tokuyasu-sectioning based Immunogold EM imaging of TMEM126A-HA labelled by anti-HA primary and 12*nm* gold secondary antibodies. White arrows indicate gold particles at cristae membranes. Scale bar: 100*nm*. E) Quantification of average distribution of immunogold particles for TMEM126A-HA for each compartment. All gold particles per mitochondria are assigned in proximity to cristae, IBM or marked as ambigious, respectively. *N* = 11 different crops of mitochondria used for quantification. The mean with the SEM is shown.

### Simulations predict experimentally validated IMM protein distributions

Here, IMM proteins with a high ratio of cristae interactions, such as NDUFAF4, DMAC2 and COX11, hint at their cristae localization. NDUFAF4 is known as an assembly factor for complex I of the ETC, similar to COX11 for complex IV, while DMAC2 is part of the ATP synthase. These known assembly factors of the respiratory chain complexes are known to be associated within the CM. On the other end, proteins with a low ratio of cristae interactions, such as TIMM8B and TIMM9 as components of the TIM complex and SLC25A1 and SLC25A11 as part of the solute carrier family, are known proteins of the IBM. ^[39–42]^ Besides the known proteins, we identified several unannotated proteins. Two proteins in the IMM were considered in particular for a detailed localization study based on the MD results as visualized in Fig. 4 C-D:

Noticeably, the two protein types considered below show less clustered neighborhoods with the IBM; proteins (Fig. 4C) compared to the cristae proteins (Fig. 4D).

The first protein ‘Family with Sequence Similarity 162 Member A (FAM162A)’ (shown in blue) is described as a mitochondrial protein with unknown function. It is speculated to be involved in apoptotic processes and having a role in maintaining mitochondrial membrane integrity. According to MitoCarta 3.0, it is localized to the IMM. With an extensive interaction landscape to cristae membrane proteins, FAM162A has a high cristae / IBM ratio, hinting at a potential cristae localization, which is also shown in Fig. 4C-D. To verify this, we performed live STED microscopy of tagged mitochondrial proteins (Halo/SNAP, HA, see also supplementary information S1) to resolve its sub-compartmental localization within the IMM using HeLa-COX8A-SNAP as a cristae reference. ^[43]^ In live STED imaging, FAM162A shows clear cristae localization, exhibiting a cristae-like striped pattern, similar to the marker COX8A (Fig. 5B).

The second protein is the ‘Transmembrane Protein 126A (TMEM126A)’ (shown in light blue). This protein is reported to be an IMM protein with a role in assembly of the mitochondrial respiratory chain complex I and in mitochondrial-encoded protein biogenesis. Recently, in Zhu *et al*. ^[17]^ and Poerschke *et al*. ^[44]^, it was shown that the known topology of TMEM126A is challenged, with the crosslinking data hinting at its N- and C-terminus facing the IMS. Furthermore, in our MD simulation based on crosslinking data, TMEM126A shows a high cristae interaction score, similar to FAM162A. As TMEM126A shows a punctate pattern in co-localization with the cristae marker in fixed cells using STED microscopy (Fig. 5C), it seems to localize rather to the cristae than the IBM. Additionally, to confirm the predicted localization of TMEM126A, Immunogold EM was performed and analyzed (Fig. 5D-E). It shows that the majority of TMEM126A is indeed found in the cristae. It has been recently reported, that TMEM126A facilitates together with OXA1L the insertion of mitochondrial-encoded proteins into the inner membrane. ^[44]^ Among these newly synthesized proteins, TMEM126A is essential for complex 1 assembly, acting as a chaperone. Our results suggest that TMEM126A acts within the cristae, facilitating insertation at the IBM in proximity to OXA1L and facilitating lateral distribution into the cristae.

## Conclusion

Cross-Linking Mass Spectrometry (XL-MS) can be used to characterize the whole interactome of cells and organelles, or, more specifically mitochondria. However, the resulting interaction networks are dense and convoluted. A straight forward interpretation through, e.g., clustering is not possible. Here, we presented a molecular dynamics (MD) model, which is directly informed by XL-MS, quantitative mass spectrometry data with super-resolution microscopy and 3D volumetric imaging. The resulting simulations revealed more nuanced clusters and interaction pattern, which allowed the identification of known and unknown protein complexes within mitochondria.

Therefore, the MD simulations allowed the qualitative interpretation of protein clusters in soluble compartments, such as the PDH complex in the matrix compartment, or the ATP-5 synthase in the inner mitochondrial membrane.

Our results suggest that the FAM162A and TMEM126A proteins are localized in the cristae, further illuminating the exact locations of proteins and expanding the mitochondrial location carta. Whereas Zhu *et al*. exclusively used the CLASP approach for localization annotation, the integration of the different methodologies presented here, enabled an increased resolution and a direct view of protein neighborhoods. Hence, the combination of computational and experimental strategies provided insights into how the proteins distributed in their respective compartments.

Despite its noted capacity, it is important to acknowledge that the model is highly simplified and does not fully capture the complexity of the system. Specifically, due to insufficient information, proteins are represented as spherical beads, which omits specific and orientational interactions. This limitation could potentially be addressed by modeling proteins with multiple beads. ^[46–49]^ However, calibrating such a system would require additional experimental data from a broader range of techniques. Coupling the model with a machine learning-driven force field might provide a solution to these challenge. ^[50–52]^ Furthermore, as the XL-MS data directly informs the interactions, a more comprehensive mitochondrial interactome with extensive quantitative crosslinking data could significantly enhance the model. Introducing novel crosslinkers could capture interactions beyond lysine proximities, extending to all amino acids, and potentially revealing interactions in lysine-deficient membrane proteins and low-abundance proteins. However, with the increase in the quality of the XL-MS data, the secondary data needs to improve as well, i.e., abundance, localization, volume, protein size.

It is worth noticing that, the data and databases used were obtained by bottom-up proteomics that cannot distinguish proteoforms. ^[53]^ An alternative could be top-down proteomics; however, these approaches are not fully feasible due to sample complexity and standardization.^[54]^

In summary, we presented an integrative molecular dynamics approach to extract valuable information from cross-linking mass spectrometry data and combining different experimental approaches. In addition to its application in analyzing XL-MS data, our model provides proof-of-concept results demonstrating that XL-MS is a valuable source of moldeing calibration data. This approach is highly beneficial for calibrating mesoscale and coarse-grained modeling techniques, potentially paving a new path for the emerging frontier of biomodeling, particularly in whole-cell and organelle-level modeling endeavors.

## Supplementary Information

**S1:** Experimental details

- Crosslinking MS
- Mitochondrial compartment localization information
- Protein abundance estimation
- 3D volumetric data
- Radii of gyration
- STED imaging

**S2:** Simulation details molecular dynamics

**S3:** Calibration of the interaction strength

**S4:** Table of neighbors over all replica simulations (.csv)

**S5:** Reduced Matrices from S4 for respiratory complexes

**S6:** Table of standard deviations of neighbors over all replica simulations (.csv)

### Acknowledgements

We thank Marie Bieck and Rozemarijn Eva van der Veen for their help in providing a more independent counting analysis of the immunogold experiments. We thank Martina Ringling and Svea Hohensee for assistance with ultrathin cryosectioning, immunogold labeling and TEM imaging. We thank Johannes Broichhagen for generously providing STED dyes for SNAP and Halo labeling.

W. P. acknowledges funding from the Novo Nordisk Foundation (grant No. NNF18SA0035142) and Marie Skłodowska-Curie Fellowship (grant No. 101104867). This research is supported by the Novo Nordisk Foundation (grant No. NNF22OC0079182) and Independent Research Fund Denmark (grant No. 10.46540/2064-00032B).

The work was funded by Deutsche Forschungsgemeinschaft Grant (DFG) SFB 958(Z03), DFG Project LI 3260/5-1, DFG Project LI 3260/6-1, the Leibniz-Wettbewerb (K284 /2019 and P70/2018), and ERC-2020-StG (project number 949184) to F.L. as well as an FMP integrative project to M.L. and F.L.

## Conflict of Interest

F.L. is a shareholder and advisory board member of Absea Biotechnology Ltd. and VantAI. The remaining authors declare no competing interests.

## Twitter Handles

- Fabian Schuhmann: @FabianSchuhmann
- Fan Liu: @theliulab
- Weria Pezeshkian: @WPezeshkian

## Notes

### Summary of Updates

The article has undergone a review process and the updated version now addresses the comments raised.

