## Supplementary material for "Integrative Molecular Dynamics Simulations Untangle Cross-Linking Data to Unveil Mitochondrial Protein Distributions": S5

### Protein identities of respiratory chain clusters

A) Cluster I

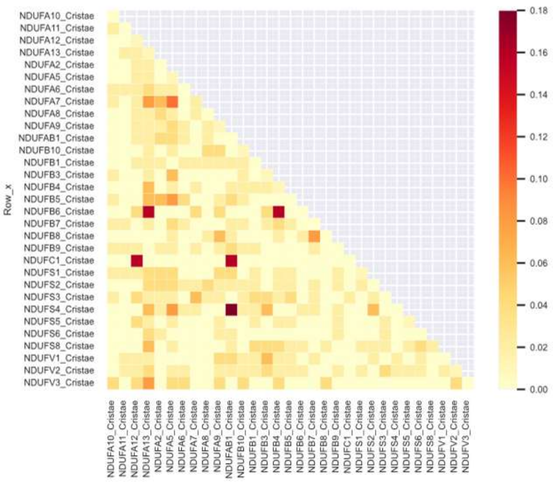


B) Cluster II

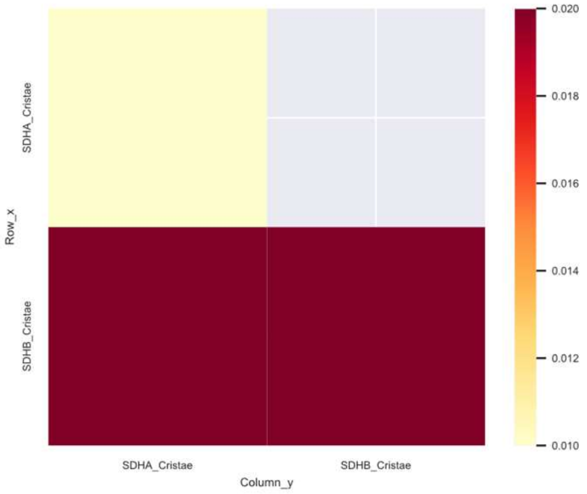


C) Cluster III

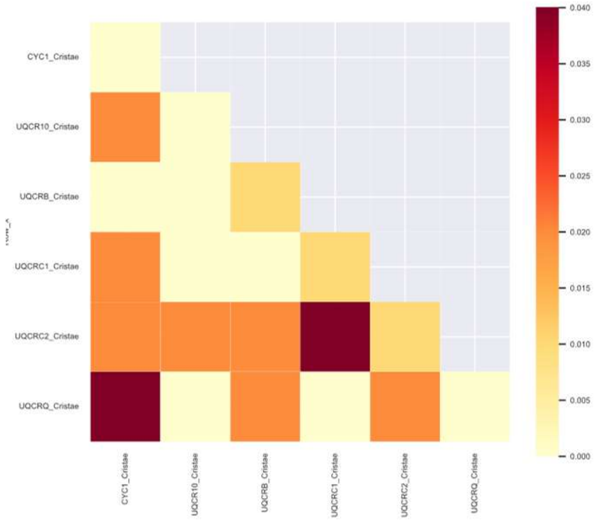


D) Cluster IV

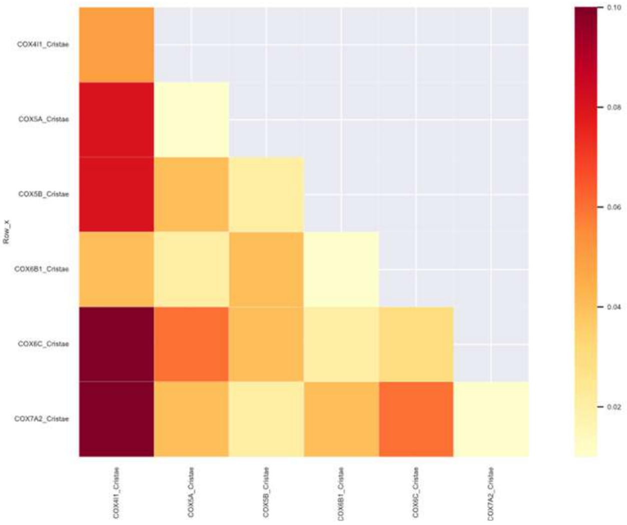


E) Cluster V

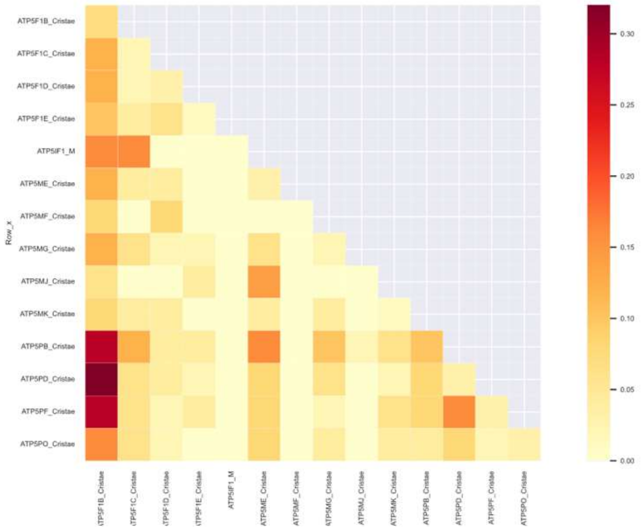


A – E)
Comparison of the MD simulation interaction score for the identified subunits of the respective clusters I – V. Complex association is based on data retrieved from HGNC.
