## Supplementary material for "Integrative Molecular Dynamics Simulations Untangle Cross-Linking Data to Unveil Mitochondrial Protein Distributions": S1-S3

#### S1: Experimental Details

##### Crosslinking MS

The crosslinking data is obtained from Zhu *et al.*[1] publication, with the dataset generated as described in the methods section. In short, crosslinking experiments were performed with samples at up to 8 mg/mL, using 0.5 mM azide-tagged acid-cleavable disuccinimidyl bissulfoxide (DSBSO) in dimethyl sulfoxide (DMSO), incubated for 30 minutes at room temperature and quenched with 30 mM TrisHCL buffer. Samples were centrifuged, washed and prepared for mass spectrometry (MS) analysis by denaturation with lysis buffer, reduction with dithiothreitol (DTT) and alkylation with chloroacetamide (CAA). Proteins were digested with Lys-C and trypsin, followed by peptide desalting using Sep-Pak C18 cartridges and enrichment using dibenzocyclooctyne (DBCO) beads. The enriched peptides underwent size exclusion chromatography (SEC) and high pH chromatography fractionation, followed by LC-MS/MS analysis using an UltiMate 3000 RSLC nano LC system and an Orbitrap Fusion Lumos mass spectrometer. MS data were analyzed with XlinkX v2.0 and results were filtered to 2% FDR at the crosslink level and protein-protein interaction networks were visualized with Cytoscape.

[a] F. Schuhmann, W. Pezeshkian
Niels Bohr Institute
University of Copenhagen
Blegdamsvej 17
2100 Copenhagen, DenmarkInstitute and organisation address



[b] K. C. Akkaya, F. Liu
Department of Structural Biology
Leibniz-Forschungsinstitut für Molekulare Pharmakologie (FMP)
Robert-Roessle-Str. 10 13125 Berlin, Germany

[c] K. C. Akkaya, D. Puchkov, M. Lehmann
Department of Molecular Physiology and Cell Biology
Leibniz-Forschungsinstitut für Molekulare Pharmakologie (FMP)
Robert-Roessle-Str. 10 13125 Berlin, Germany

[d] F.Liu
Charité – Universitätsmedizin Berlin
Charitépl. 1, 10117, Berlin, Germany

[+] These authors contributed equally.

##### Mitochondrial compartment localization information

Data on sub-mitochondrial localization of proteins is based on an earlier study by Rath *et al.*[2]. Additional sub-compartment localization for distinguishing markers of IBM and cristae membrane is extracted based on proteins and their complex-association from HGNC to the main complexes of the ETC, TIM-complexes and MICOS-complex.

##### Protein abundance estimation

The mitochondrial quantitative copy numbers are obtained from Morgenstern *et al.*[3] and were generated as described in the method section. In short: Morgenstern *et al.* estimates the mitochondrial protein copy number based on MS1 intensities determined from whole cell lysates. This approach uses a combination of the total protein approach (TPA) and line fitting the resulting copy numbers per cell as a proteomic ruler to acquired mitochondrial iBAQ measurements of bona fide proteins. The TPA method is a label- and standard free-method for absolute protein quantification, based on spectral intensities acquired in the large-scale proteomic analyses and can be extended to allow the calculation of protein abundances.

##### 3D volumetric data

To determine the relative mitochondrial volume fraction versus the cytoplasmic volume in HEK293T cells, the cells were seeded on 8-well ibidi™ chambers and, the following day, stained with 0.1 $\mu$M MitoTracker DeepRed FM for mitochondria, 5 $\mu$M CellTracker Green CMFDA for cytoplasm, and 1 $\mu$M pHoechst for nuclei. After incubating for one hour at 37°C, the cells underwent extensive washing and a one-hour post-incubation for dye removal, then imaged in live-cell microscopy with live-cell imaging buffer and 10% FBS. Z-stacks of single cells were acquired using a confocal spinning disc microscope with a 0.3 $\mu$m step size, covering the entire cell volume. The z-stacks were analyzed in Imaris software to create 3D models of the cell and mitochondria, from which volumetric measurements were extracted. As an orthogonal control, measurements were performed also at 2D (SEM) and 3D (FIB-SEM) electon microsopy images. Volume fraction of mitochondria in cytoplasm, mitochondrial matrix volume, relative area of OMM, IMM and cristae were quantified by stereology methods for volume fraction and surface density estimation.[4] For volumetric analysis of individual mitochondria, 3D cellular volume was acquired by FIB-SEM and 3D segmentation of individual mitochondria was performed by using deep learning workflow of Microscopy Image Browser.[5] Mitochondrial networks were manually refined and 3D model was generated in IMARIS. In total, using confocal microscopy of living cells, the total volume of mitochondria was measured at an average of 445 $\mu m^3$, representing roughly 8.6% of the total cellular volume. In contrast, using volumetric EM based on focused ion beam scanning electron microscopy (FIB-SEM), we measured a relative mitochondrial volume fraction of 12%. With this, we used it to calculate a rough absolute mitochondrial volume based on the total HEK293T volume acquired in the confocal microscopy approach ($\sim 5,400\ \mu m^3$), resulting in roughly 640 $\mu m^3$ per cell. Combining both approaches estimates the average mitochondrial volume to 543 $\mu m^3$ per HEK293T cell.

#### Radii of gyration

For any given mitochondrial protein in the crosslinking dataset, the radius of gyration, as a globular representation of the protein, was estimated using the structure generated by AlphaFold2 and a Python-wrapped version of the "rgyr" function from Bio3d.

#### STED imaging

HeLa cells and HeLa-COX8A-SNAP cells were seeded in $\mu$-Slide 8-well ibidi$^{TM}$ chamber (80827) and cultured in DMEM with 10% FBS, 1% P/S, and 1% L-glutamine. For SNAP-tag labeling, cells were incubated for one hour with 1 $\mu$M BG-SiR-d12 in DMEM at 37°C, followed by three washes with DMEM before imaging.[6] For Halo-tag labeling, 1 $\mu$M of CA-JF542 or CA-SiR-d12 was used. Live cell microscopy was performed in HEPES Imaging buffer with 10% FBS at 37°C or in DMEM with CO2 support. Cells to be fixed were stained before fixation. Cells were fixed with PBS containing 4% PFA and 4% sucrose for 10 minutes at RT. For preserving mitochondrial ultrastructure, cells were fixed in-culture with 37°C prewarmed fixation solution for 20 minutes. Fixation was quenched with PBS containing 0.1 M glycine and 0.1 M ammonium chloride for 10 minutes at RT, permeabilized with PBS containing 0.15% TritonX-100 for 10 minutes, and washed with PBS. Cells were blocked with PBS containing 1% BSA and 6% NGS for 30 minutes at RT, incubated with primary antibodies in blocking buffer for one hour at RT or overnight at 4°C, washed, and then incubated with secondary antibodies for 30 minutes at RT. After washing, cells were directly imaged. STED microscopy was performed using a STEDYCON system on a Nikon TI Eclipse microscope, with live cell imaging at 37°C or fixed cell imaging at RT. The STEDYCON system used 561 nm and 640 nm lasers for excitation, paired with avalanche photodiodes for detection, a 775 nm STED laser for depletion, and an optical tunable filter for confocal imaging. Images were captured using a 100x oil objective with a 1.45 NA, a time gate of 0.5–8 ns, 8x line averaging, and a final pixel size of 20 nm for STED images.

#### Immunogold EM and quantification

Cells were fixed by 4% PFA in PBS, scrapped from the dish and centrifugated into the pellet at 2000g, embedded into 2% agarose (PBS). Pellets were cut into $0,5-1\ mm^3$ pieces and further processed for ultrathin cryosectioning and immunogold labeling as in Overhoff *et al.*[7]. For staining, 1:50 Rabbit Anti HA (Caymann #94) and 1:50 12 nm gold goat anti-rabbit antibodies (Dianova) were used. Staining of cells not expressing TMEM126A-HA has not revealed any detectable immunogold labelling in comparable sized observed area of cellular profiles.

For relative quantification of the localization of TMEM-126A-HA Immunogold particles in different subcompartments, mitochondria were manually segmented within broad overviews. Subsequent classification into IBM, Cristae, and Matrix was performed individually by 3 individuals depending on the localization of Gold particles in close proximity to parallel membranes (Cristae), single orthogonal membranes (IBM), and ambiguous localization (Matrix).

### S2: Molecular Dynamics

In order to derive the simulation framework and parameters, experimental and computational approaches were combined. For a given protein, the radius of gyration was estimated using the structure generated by AlphaFold2[8] and its abundance was quantified through the experimental method. Furthermore, for each pair of protein types, the number of crosslinks between them was taken from the experimental data. These data were then translated to a Lennard-Jones potential interacting between protein $A$ and protein $B$:

$$LJ(A,B) = 4\varepsilon(A,B)\left(\left(\frac{\sigma(A,B)}{r(A,B)}\right)^{12} - \left(\frac{\sigma(A,B)}{r(A,B)}\right)^{6}\right), \tag{1}$$

with $\sigma(A,B)$ denoting the averaged radius of gyration of proteins $A$ and $B$, $r(A,B)$ is the distance between them and $\varepsilon$ is given by

$$\varepsilon(A,B) = \rho\frac{XL_{A,B}}{N_A + N_B}. \tag{2}$$

Here, $XL_{A,B}$ is the number of crosslinks between proteins $A$ and $B$; $N_A$, $N_B$ denote the abundance of the protein type. The interactions are scaled by the parameter $\rho$, which is set and discussed in the calibration S3: Calibration of the interaction strength.

Each protein is represented by a single bead, which is randomly placed into layers, which correspond to the location of the protein in mitochondria. For the simulation, each layer is 500 × 500 nm in the surface. Membranes are represented in two-dimensions, while non-membrane proteins inhabit a space with a height of 14 nm. The height for each compartment is introduced using a flat-bottom potential goverend by a flat-bottom potential between the membranes. A flat-bottom potential will not interfere with the motion of the protein, unless it attempts to leave its confined space and penetrate a membrane[9]. An empty virtual membrane keeps the IMS not bounded by membranes from moving away from the IMM/Cristae membrane.

For protein regions, which appear more than once, the number of proteins was multiplied and the compartment was chosen randomly with equal weights equally distributing the proteins.

The whole system was subsequently simulated using GROMACS[9–16]. The system is minimized using a steepest descent algorithm. Soft core minimization is employed to allow the minimization of potentially overlapping proteins in the random placement. The simulation of the system is conducted for 230.4 $\mu$s with an integration time step of 0.2 ps and the leap-frog stochastic integrator. The system is semi-isotropically coupled with a temperature of 310 K.

The simulation was repeated with different initial random placement 30 times.

### S3: Calibration of the interaction strength

The interactions have to be calibrated, and thus the factor $\rho$ is introduced to the Lennard-Jones potential described in S2. In order to estimate the magnitude of $\rho$ test simulations were conducted only containing the OMM membrane, as it contains a variety of known complexes. In these test simulations, $\rho$ was chosen to be between 10 and 90 with a step size of 10. For each step, ten replica simulations were performed. All other parameters are equal to the ones discussed in S2. The effect of $\rho$ is visualized in Fig. SF1.

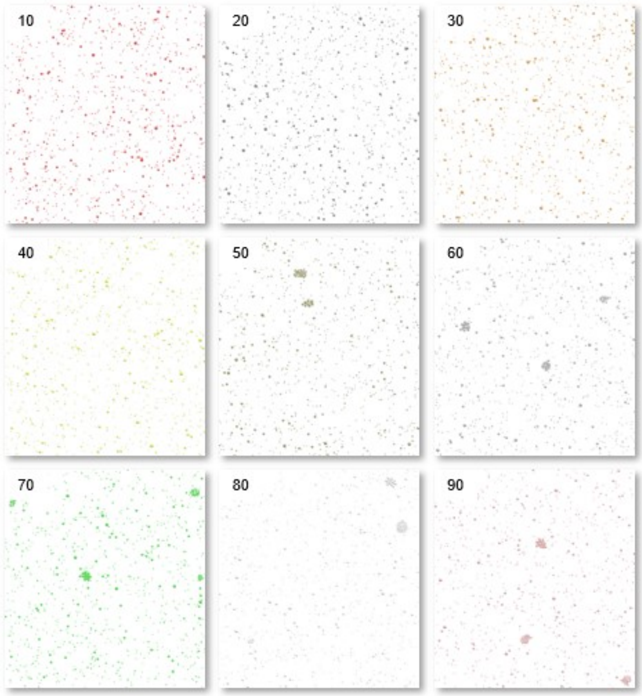


**Figure SF1.** A top view of the OMM membrane used for calibration is shown for different values of $\rho$ from the top left at $\rho = 10$ to the bottom right at $\rho = 90$. At $\rho = 50$, first clusters appear, while the proteins remain unclustered for lower $\rho$.

Formed clusters were quantitatively analyzed using a hierarchical clustering approach. Based on the last frame of each simulation, the distances between all the proteins were computed and clustered. For the clustering a distance threshold of $24.29 \cdot 2 \cdot 1.25$ has been chosen, which corresponds to the average diameter of the proteins based on their radius of gyration times an $\varepsilon$ to allow the clustering of larger than average proteins. Figure SF3 shows the average number of clusters with at least three proteins for each value of $\rho = \{10, 20, \cdots, 90\}$.

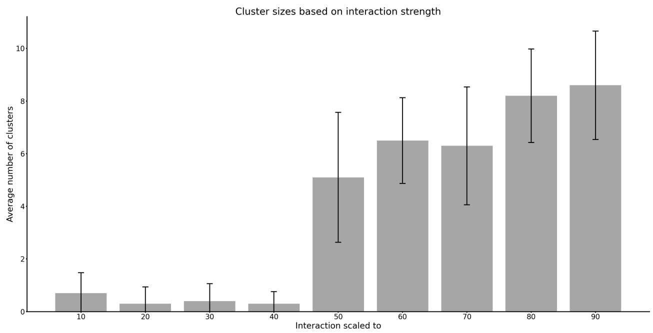


**Figure SF2.** The average number of clusters per interaction strength through the replica is shown including the standard deviation as error bars. Based on the data, $\rho = 60$ was chosen as it shown consistently decent cluster formation with a low deviation.

[8] J. Jumper, R. Evans, A. Pritzel, T. Green, M. Figurnov, O. Ronneberger, K. Tunyasuvunakool, R. Bates, A. Žídek, A. Potapenko, A. Bridgland, C. Meyer, S. A. Kohl, A. J. Ballard, A. Cowie, B. Romera-Paredes, S. Nikolov, R. Jain, J. Adler, T. Back, S. Petersen, D. Reiman, E. Clancy, M. Zielinski, M. Steinegger, M. Pacholska, T. Berghammer, S. Bodenstein,

D. Silver, O. Vinyals, A. W. Senior, K. Kavukcuoglu, P. Kohli, D. Hassabis, *Nature* **2021**, *596*, 583.
